# Increase in stable isotope ratios driven by metabolic alterations in amphipods exposed to the beta-blocker propranolol

**DOI:** 10.1101/520205

**Authors:** Caroline Ek, Zhenyang Yu, Andrius Garbaras, Hanna Oskarsson, Ann-Kristin Eriksson Wiklund, Linda Kumblad, Elena Gorokhova

## Abstract

Anthropogenic pressures, such as contaminant exposure, may affect stable isotope ratios in biota. These changes are driven by alterations in the nutrient allocation and metabolic pathways induced by specific stressors. In a controlled microcosm study with the amphipod *Gammarus* spp., we studied effects of the β-blocker propranolol on stable isotope signatures (δ^15^N and δ^13^C), elemental composition (%C and %N), and growth (protein content and body size) as well as biomarkers of oxidative status (antioxidant capacity, ORAC; lipid peroxidation, TBARS) and neurological activity (acetylcholinesterase, AChE). Based on the known effects of propranolol exposure on cellular functions, i.e., its mode of action (MOA), we expected to observe a lower scope for growth, accompanied by a decrease in protein deposition, oxidative processes and AChE inhibition, with a resulting increase in the isotopic signatures. The observed responses supported most of these predictions. In particular, %N was positively affected by propranolol, whereas both protein allocation and body size declined. Moreover, both ORAC and TBARS levels decreased with increasing propranolol concentration, with the decrease being more pronounced for TBARS, which indicates the prevalence of the antioxidative processes. These changes resulted in a significant increase of the δ^15^N and δ^13^C values in the propranolol-exposed animals compared to the control. These findings suggest that MOA of β-blockers may be used to predict sublethal effects in nontarget species, including inhibited AChE activity, improved oxidative balance, and elevated stable isotope ratios. The latter also indicates that metabolism-driven responses to environmental contaminants can alter stable isotope signatures, which should be taken into account when interpreting trophic interactions in the food webs.

## Introduction

In human physiology, the natural variations of the isotopic ratios of carbon, nitrogen and other major elements comprising biomass (δ^15^N, δ^13^C, δ^18^O, δ^2^H, δ^44/40^Ca, etc.) are attracting increasing attention since they offer a new means to study the imbalances linked to pathological conditions [1,2]. In non-human biology, however, the physiology of a consumer is rarely coupled to δ^15^N and δ^13^C values that are assumed to be a bare reflection of the diets’ isotope composition plus a discrimination factor (Δ^15^N and Δ^13^C, respectively). There is ample evidence that consumer Δ-values and the isotopic signatures may vary depending on various endogenous and environmental factors via their effects on metabolism and growth. These factors include variations in moulting status [3], food quantity [4,5] and quality [6], temperature [7,8], and contaminant exposure [9]. Therefore, to improve interpretation of stable isotope analysis (SIA) data in stress ecology and ecotoxicology, it is crucial to consider the physiological state in addition to the potential dietary sources of the consumer.

In line with this, isotope signatures have been reported to respond to changes in oxidative status [10], suggesting that both δ^15^N and δ^13^C values can reflect not only the diet but also shifts in the balance between antioxidative and pro-oxidative processes. Such shifts are not particularly unexpected because heavier isotopes have been found to accumulate in molecules in which they are present in the highest oxidation state [11]. Oxidative stress is commonly used to assess toxic exposure [12] and various metabolic activities related to oxidative balance, such as feeding [13] and reproduction [14]. Therefore, by combining biomarkers and SIA data, the assessment of trophic structure based on SIA can be facilitated by including responses to confounding factors, such as toxic exposure, migratory activity, and food levels. It has also been proposed that major biomolecules (i.e., proteins, nucleic acids, and lipids) enriched in heavy isotopes might be less prone to the oxidative damage due to kinetic effects and higher stability [15,16]. Hence, organisms and tissues with higher isotope signatures would display lower oxidative damage. On the other hand, under toxic exposure variations in δ-values may be explained by changes in the relative abundance of heavy and light isotopes due to the elevated oxidation.

Studies exploring effects of contaminants on δ^15^N and δ^13^C values have largely focused on POPs, persistent organic pollutants [8,9,17,18]. POPs are known to cause deleterious effects in wildlife via oxidative stress, increased physiological costs, and neurotoxicity [12]. A group of very different environmental contaminants is pharmaceuticals. First, they are designed to be biologically active in living organisms by having a specific drug target, which is a molecular structure that undergoes a specific interaction, i.e., the mechanism of action (MOA), with the drug administered to treat or diagnose a disease. MOA connects specific molecular and metabolic interactions to the response, but for most drugs, several (if not many) targets are known. Second, the potency of a pharmaceutical for a non-target organism is dependent on whether drug targets are evolutionary conserved [19], and unknown targets and MOAs for such drugs cannot be excluded in phylogenetically distant species. Whereas the effects of toxic exposure on stable isotope ratios are often associated with increased physiological costs due to detoxification, such effects of pharmaceuticals may result from the intended therapeutic effects or side effects. Hence, to understand changes in the isotopic composition of consumers exposed to pharmaceuticals present in the environment, the specific MOAs need to be considered.

Propranolol is a non-selective β-blocker used to treat hypertension by acting as an antagonist to the adrenergic β-receptors. In mammals, in addition to lowering blood pressure and heart rate, propranolol has been reported to lower protein turnover via modulation of both protein synthesis and catabolism [20,21]. Furthermore, it also possesses antioxidant properties by stabilizing lysosomes [22] and, thus, reducing oxidative stress [23]. For crustaceans, it appears as there exist no adrenergic receptors in the cardiovascular system [24], which would indicate low potency for a propranolol effect. However, propranolol is also a serotonin receptor (5-HT) antagonist [25] for which there are evolutionarily conserved receptors in crustaceans and other invertebrates [26,27]. In bivalves, for example, propranolol exposure has been linked to altered cyclic adenosine monophosphate (cAMP) signaling via the 5-HT1 receptor [28], suggesting that effects on cAMP signaling in invertebrates, including crustaceans, can occur and cause downstream alterations in catabolic, anabolic and transport processes. Moreover, because of the overlapping adrenergic and cholinergic innervation in many systems, drugs acting on one system are known to modify the activity of the other. This is also the case with propranolol, which inhibits cholinesterase (ChE) enzyme activity [29]. In line with these MOAs, propranolol has been found to reduce heart rate in *Daphnia magna* [30] and motility in *Gammarus* spp. [31]. Therefore, propranolol presents a suitable model substance to predict effects of pharmaceuticals on isotope ratios in crustaceans and to test the relationships between oxidative stress and isotope ratios, as a consequence of contaminant-induced changes in the physiology.

This study aimed to evaluate whether MOAs of propranolol can be used to predict its effects on stable isotope ratios and oxidative stress biomarkers in crustaceans (Table 1; Fig. 1). We assayed elemental composition (percentages of carbon and nitrogen, %C and %N, respectively), stable isotope ratios (δ^15^N and δ^13^C) and biomarkers of oxidative stress and neurological damage in amphipods *(Gammarus* spp.) exposed to propranolol (100 and 1000 μg L^-1^) in microcosms. These test concentrations are high compared to the ecologically relevant concentrations, which have been reported to be in the upper range of 0.29-1.9 and 0.59 μg L^-1^ for sewage treatment plant effluent and surface waters, respectively [32,33]. However, using high concentrations was considered appropriate, because we were not focused on assessing the effects of environmentally relevant concentrations of propranolol in non-target species but investigated the overall predictably of these effects.

**Figure 1.**
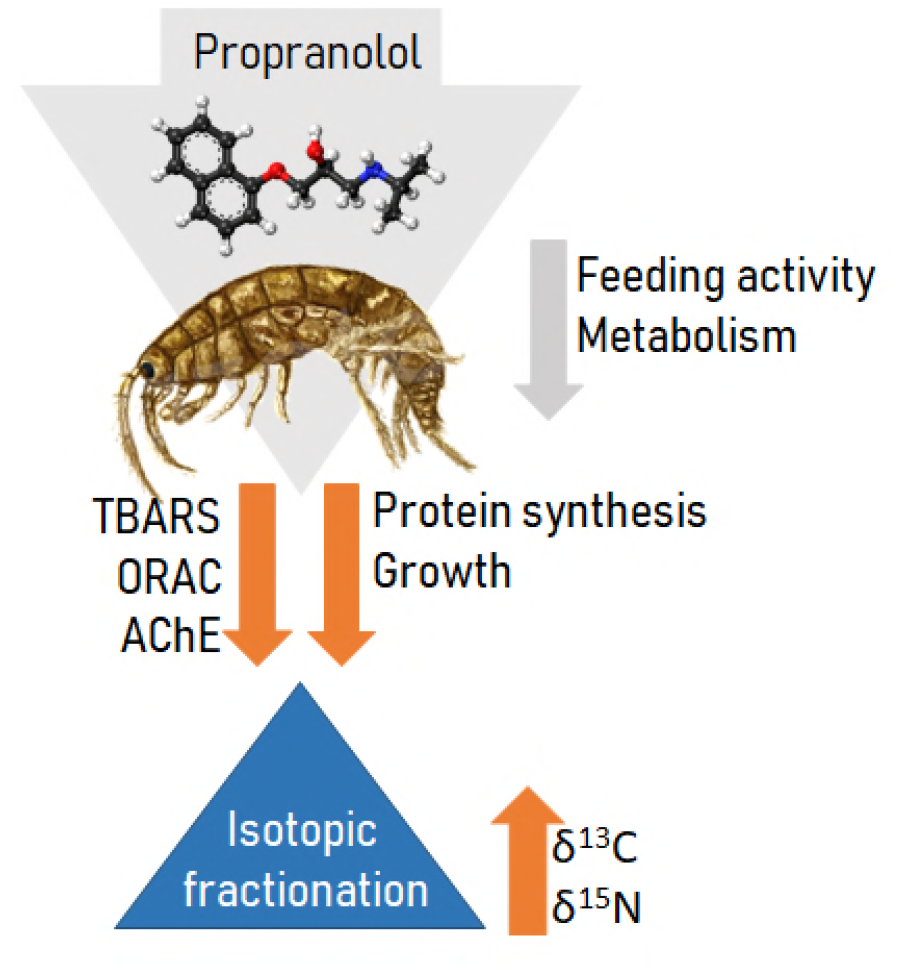
Conceptual diagram for propranolol effects on amphipod growth and its constituents, biomarkers (TBARS, ORAC, and AChE) and stable isotope signatures (δ^13^C and δ^15^N), predicted on the basis of MOA. The negative effects are shown as arrows pointing down and positive as arrows pointing up. The orange arrows depict measured effects and grey arrows indicate those that were not accessed but appear plausible. See Table 1 for the rationale for specific effects.

**Table 1.** Summary of the hypothesized relationships, the rationale behind each hypothesis, and models used in the hypothesis testing. Propranolol (μg L^-1^); %N, nitrogen content; %C, carbon content, WW, wet weight; ORAC, oxygen radical absorbance capacity as proxy for antioxidative capacity; AChE, acetylcholinesterase activity; ORAC:TBARS ratio, a proxy for the balance between antioxidative and pro-oxidative activities. Significant predictors are in bold face.

| Hypothesis | Response | Predictors | Changes expected to occur in the propranolol-exposed animals and the rationale |
| --- | --- | --- | --- |
| H1 | %N | **propranolol** | %N is positively affected by propranolol due to decreased protein catabolism, amino acid release and breakdown [21], whereas protein allocation is reduced due to negative effects of propranolol on food intake [49].<br>%N is a plausible predictor for protein content due to stoichiometric principles [45]. |
| H2 | Protein | **propranolol**<br>**%N** |  |
| H3 | WW | Propranolol<br>**Protein** | Somatic growth decreases due to inhibitory effect of propranolol on cell proliferation and metabolism, with concomitant effects on oxygen supply and energy balance [30]. |
| H4 | TBARS | **Propranolol**<br>WW | Oxidative damage decreases due to the antioxidative properties of propranolol [22] but also as a result of the reduced feeding [13]. As many other biomarkers in crustaceans are affected by size/age [54], WW may also contribute to the variability of this biomarker. |
| H5 | ORAC | TBARS<br>**WW** | Antioxidant capacity decreases due to the decreased pro-oxidative processes (see H4). WW is an ontogenetic covariate. |
| H6 | AChE | **Propranolol**<br>**ORAC:TBARS** | AChE inhibition occurs due to the propranolol binding to the peripheral sites on the AChE molecule, thus reducing its activity [49]. Also, lower levels of oxidative stress are likely to lower AChE levels as moderate oxidative stress can stimulate AChE activity [56]. |
| H7 | $\delta^{15}\text{N}$ | **Propranolol**<br>WW<br>**ORAC:TBARS** | $\delta^{15}\text{N}$ and $\delta^{13}\text{C}$ values increase in response to metabolic alterations induced by propranolol exposure. Alternatively, strictly due to the growth inhibition (H3) and concomitant increase in $^{15}\text{N}$ fractionation [5]. The increase in $^{13}\text{C}$ fractionation is related to higher mass-specific respiration rate in the exposed animals. Also, as lipids are depleted in $^{13}\text{C}$, the covariates are %C and C:N as proxies for the lipid content. Moreover, TBARS and ORAC:TBARS ratio are included as covariates reflecting oxidative status, because fatty acids with higher proportion of heavier isotopes would be less prone to oxidation [16]. |
| H8 | $\delta^{13}\text{C}$ | **Propranolol**<br>WW<br>C:N or %C<br>ORAC:TBARS | |

## Material and Methods

### Microcosms

The model communities consisting of amphipods *(Gammarus* spp.), blue mussels *(Mytilus edulis trossulus),* red filamentous macroalgae *(Ceramium tenuicorne),* were exposed to 0, 100 and 1000 μg L^-1^ propranolol in 8-L aquaria for six weeks, five aquaria per treatment. Animals and sediment used in the microcosms were collected in June 2011 nearby Askö laboratory, in the northern Baltic proper, Sweden. Details on sampling, sediment collection and quality are presented elsewhere [34]. In each of the 15 aquaria, 30 amphipods, 31 mussels, and 8.3 g WW *Ceramium* were placed. Before the experiment, microcosms were acclimated for seven days in a climate chamber with a water temperature of 12.5 ± 0.5 °C and a 16:8 h light:dark regime; the same conditions were used during the experiment. The microcosms were connected to a flow-through system with seawater (salinity 6.5 PSU) entering each aquarium via a PVC tubing allowing for the daily exchange of the total water volume. Propranolol was dissolved in dilute phosphoric acid (pH adjusted to 7.1 using sodium carbonate) and continuously added at the test concentrations via siphons of PTFE Teflon tubing connected to fused silica capillaries from glass reservoirs above the aquaria into each experimental unit. The microalgae *(Isochrysis galbana,* Reed Mariculture) were added daily *ad libitum* (7.8 × 10^7^ cells L^-1^) to ensure a sufficient food supply for the animals. The mortality was recorded daily, and dead animals were removed. Samples for quantification of propranolol in water were taken at day 2 and day15, and samples for quantification of propranolol in organisms and sediment were taken at the end of the experiment. Upon termination of the experiment, the wet weight (WW) of the surviving amphipods was determined, and they were frozen individually in Eppendorf tubes at -80 °C pending biomarker and stable isotope analysis. The general responses to the exposure (i.e., mortality, respiration, feeding rate, excretion and community-specific gross production and respiration) as well as the concentrations of propranolol quantified for the different compartments are reported elsewhere [34]; mortality and propranolol quantifications for water and amphipods reported in this publication and relevant for our results are summarized in Supporting Information, Table S1. Here, we focus on the propranolol effects on the stable isotope signatures, elemental composition and biomarkers in the amphipods collected within the same experiment as well as the relationships between these endpoints.

### Biomarker selection

Assessment of oxidative status was conducted using total oxygen radical absorbance capacity (ORAC) and lipid peroxidation (TBARS) assays, whereas acetylcholinesterase (AChE) activity was used to evaluate effects of propranolol on AChE inhibition; see Table 1 for details on the expected effects and causal relationships between the endpoints and biomarkers. The antioxidant capacity (ORAC) reflects concentrations of water-soluble antioxidants, the substances that delay or prevent the oxidation of biomolecules by reactive oxygen species, ROS [35]. When pro-oxidative processes dominate, the reaction between ROS and lipids gives rise to lipid peroxidation (assayed here as TBARS) that causes functional loss of membrane-stability. Such shifts in the oxidative status have been linked to several diseases and aging [36]. Also, the ORAC:TBARS ratio can serve as a proxy for the balance between the antioxidative and pro-oxidative processes with lower values indicating the prevalence of oxidation [37]. The activity of AChE is central for maintaining the function of acetylcholine receptors by degrading the neurotransmitter acetylcholine; various neuropathologies, including decreased cardiac output and motility, occur in animals with AChE inhibition [38].

### Sample preparation

For biomarker and stable isotope analyses, 45 amphipods (15 ind. treatment^-1^) were used. Using a stereomicroscope and a scalpel, each individual was dissected separating abdominal and thoracic parts; during the dissection, the animals were held on dry ice. The thoracic part was used for biomarker analysis, while the abdominal part was used for δ^15^N and δ^13^C analysis. The SIA samples were placed in pre-weighed tin capsules, dried at 60 °C for 24 h, weighed and stored in a desiccator before being shipped to the SIA facilities.

### Biochemical assays

Using FastPrep (MP Biomedicals), the samples were homogenized in 500 μL of PPB (0.1 M, pH 7.2) for 30 sec × 2 times with an ice bath (30 sec) in between. After centrifugation (10 000 rpm × 5 min at 4 °C), the supernatants from the two tubes were aliquoted to 40, 50 and 150 μL for protein, ORAC, TBARS, and AChE, respectively; the aliquots were frozen at -80 °C pending the analyses. All samples were analyzed in duplicate for each biomarker using a FluoStar Optima plate reader (BMG Lab Technologies, Germany) with absorbance (protein and AChE) or fluorescence (ORAC and TBARS) configuration.

#### Protein Quantification

Protein content (μg ind.^-1^) was determined by the bicinchoninic acid (BCA) method [39] using a Pierce BCA Protein Assay kit (Thermo Scientific, Product No. 23225) and the microplate protocol with some modifications. In transparent 96-well microplate, 20 μL sample and 130 μL working solution were mixed in each well. Absorbance was measured at 540 nm, and protein concentrations were calculated using a standard curve (18.5-1500 μg mL^-1^).

#### ORAC

We used the microplate-based assay [35] with fluorescein as a fluorescent probe (106 nM), 2,2-azobis(2-amidinopropane), dihydrochloride (AAPH; 152.66 mM) as a source of peroxyl radicals, and Trolox (218 μM) as the standard. The samples were diluted with PPB (0.1 M, pH 7.2) to 0.08 – 0.12 mg protein mL^-1^ and 25 μL were mixed with 30 μL AAPH and 150 μL fluorescein in each well. After 5 min incubation at 37 °C, a kinetic fluorescence read (excitation 485 nm, emission 538 nm, 2 min/cycle × 65 cycles) was carried out. The values of the area under the curve (AUC) of standards were used to calculate the ORAC levels per individual. These ORAC values were normalized to protein content to account for variability in wet mass.

#### Lipid Peroxidation

The TBARS was measured as aldehydic lipid peroxidation fluorescent products after reacting with thiobarbituric acid (TBA) [40]. The samples were first diluted with PPB (0.1 M, pH 7.2) to 0.50 mg protein mL^-1^. Then, homogenate (150 μL) was diluted 1:1 with 10% trichloroacetic acid and rested on ice for 5 min. After centrifugation (10 000 rpm × 5 min at 4 °C), the supernatant (250 μL) was mixed with 150 μL of reaction solution (200 mg TBA in 5 mL 1.5 M NaOH and 5 mL acetic acid), and incubated at 100 °C for 1 h. After cooling to room temperature and the addition of butanol/pyridine mixture (250 μL; volume ratio 15:1), the samples were vortexed (2 × 10 s) and centrifuged for 5 min at 4000 g at 20 °C. The organic phase (upper layer, 80 μL well^-1^) was used for fluorometric determination (excitation 540 nm, emission 590 nm) of malondialdehyde (MDA) concentration (μM MDA equivalents ind.^-1^). The MDA concentrations were expressed as TBARS normalized to protein content of the sample.

#### AChE activity

Ellman assay [41] modified for a microplate format was used. The reaction solution was freshly prepared by mixing 169 μL acetylthiocholine solution (75 mM), 845 μl DTNB solution (10 mM), and 25.4 mL PPB. In each well, sample (25 μL) was mixed with 250 μL reaction solution. The plate was then incubated for 2 min with a gentle shake in the plate reader, followed by reading absorbance at 405 nm every 2 min with ten cycles. The AChE activity was calculated as:

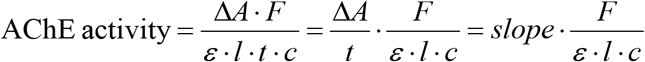

where slope is calculated through a linear fit between the absorbance change (sample minus blank) and time, *F* refers to the dilution factor (total volume/sample volume, 275 μL/25 μL), *ε* is extinction coefficient of DTNB (13600 M^-1^ cm^-1^), *l* is the length of light path length (0.75 cm in the present instrument) and *c* is the protein concentration (mg mL^-1^).

### Stable isotope ratio analysis

Stable isotope analysis was conducted at the Center for Physical Science and Technology, Vilnius, Lithuania, using a Flash EA 1112 Series Elemental Analyzer connected via a Conflo III to a DeltaV Advantage isotope ratio mass spectrometer (all Thermo Finnigan, Bremen, Germany). The stable isotope ratios ^15^N:^14^N and ^13^C:^12^C are expressed relative to the international standards atmospheric air (N) and Vienna Pee Dee Belemnite (C). Caffeine (IAEA-600) was used as secondary reference material for the reference gas calibration.

Elemental composition of nitrogen and carbon (%N and %C, respectively) are expressed as the percentage content of the sample dry weight. Calibration curves for %N and %C quantification were created using EMA P2 reference material (Elemental Microanalysis). To estimate the analytical precision of δ^15^N and δ^13^C, an internal reference *(Esox lucius,* n = 6) was analyzed together with the test samples. For analytical precision of elemental composition (%N and %C), a series of samples (n = 9) each containing a single individual of the crustacean *Daphnia magna* collected from a culture (size 340-520 μg) were used. The overall analytical precision was 0.1 ‰ and 0.04 ‰ for δ^15^N and δ^13^C, respectively, and 0.02% and 0.09% for %N and %C, respectively.

### Data analysis and statistics

To explore the overall variability in the data set, a between-group principal component analysis (bgPCA) was conducted using PAleontological STatistics (PAST) version 3.13 [42]. In the bgPCA, we used a correlation matrix of %N, %C, C:N, protein content, WW, ORAC, TBARS, ORAC:TBARS, AChE, δ^15^N and δ^13^C. When necessary, some variables (%C, WW, TBARS, ORAC:TBARS, AChE, δ^15^N and δ^13^C) were Box-Cox transformed using Statistica 8.0 (StatSoft, USA).

As the next step, all predictors were evaluated for aquarium effects, because the experimental design did not allow for complete independence between the amphipods sampled from the same experimental unit. Linear mixed models (LMM) with restricted maximum likelihood (REML) was used to test for random effect, i.e., whether the fixed effect of treatment was different between the aquaria. Linear models using generalized least square (GLS) with REML (without random effects) were used as null models to test the hypotheses of a significant aquarium effect for different variables; the resulting models were compared using analysis of variance (ANOVA). When no significant aquarium effect was detected (%N, WW, protein, δ^5^N, δ^13^C, and all biomarkers), generalized linear models (GLMs) were used to analyze the data. When this effect was significant (%C and the C:N ratio), the data were analyzed using GLS-REML. Thus, to answer the hypotheses linking specific responses to propranolol effects, we used either GLM (H1 to H6) or GLS-REML (H7; Table 1). All linear mixed models and linear models using GLS were made using the package *Linear and Nonlinear Mixed Effects Models* (nlme) version 3.1-118, and for GLMs *The R Stats Package* (stats) was used in the statistical software R version 3.1.2 (R core team 2015). The significance level was set to α ≤ 0.05. If not specified otherwise, the data are reported as mean and standard deviation.

## Results

### Principal component analysis

The bgPCA suggested a lower similarity between the animals exposed to the highest propranolol concentration (PH) and the control compared to that for the PL and control groups (Figure 2). The first principal component (PC1) described 85.8% of the variance and was best explained (>0.3) by the negative loadings of %N, ORAC:TBARS ratio, δ^15^N and δ^13^C. The PC1 also contributed most to the between-group differences, with positive loadings of C:N ratio, protein content, WW, TBARS, and ORAC. The projection of PH treatment on PC1 suggested that high propranolol exposure coincided with increased %N yet decreased protein content and lowered oxidative damage, i.e., low levels of TBARS and high levels of ORAC:TBARS ratio (Table 2).

**Figure 2.**
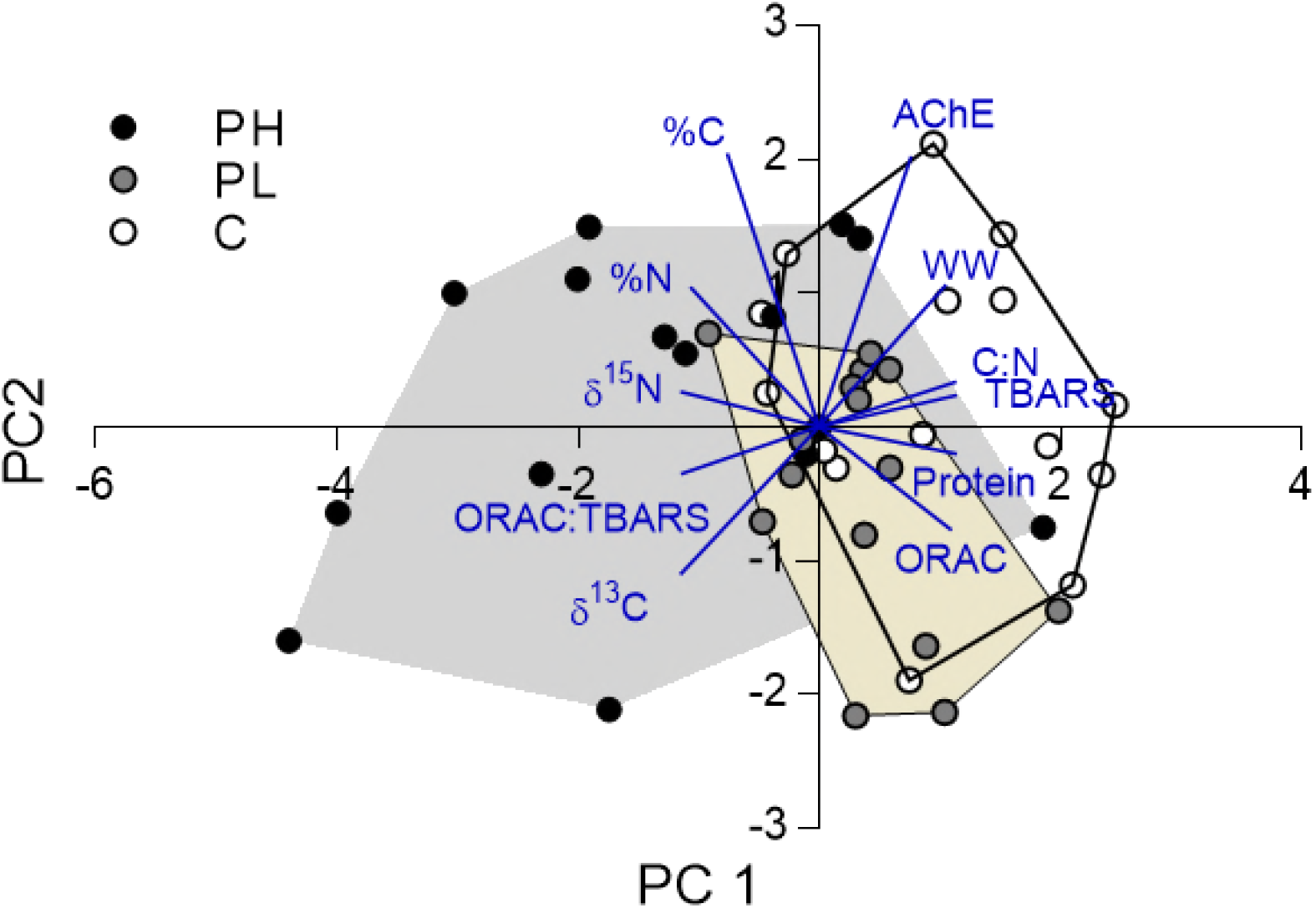
Between group principal component analysis (bgPCA) of the amphipod responses to propranolol exposure. The animals exposed to the high concentration (PH, grey) showed the least overlap with those in Control (open) and low concentration (PL, beige) exposure. PC1 and PC2 explained 85.8% and 14.2% of the variation, respectively. The vectors represent loadings for specific variables (Table 2).

**Table 2.**
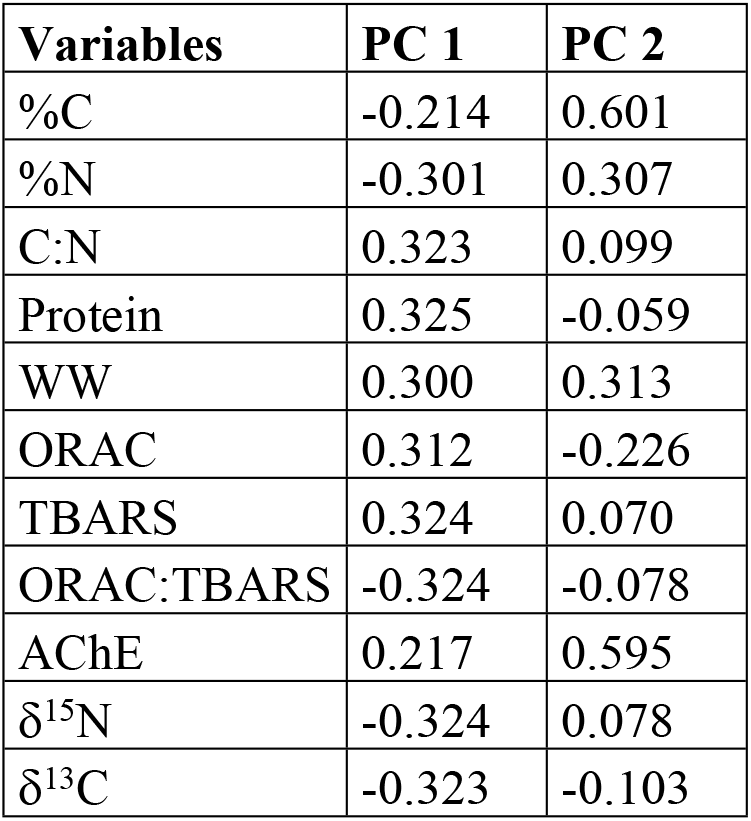
Principal component loadings for component 1 (PC1) and 2 (PC2).

| **Variables** | **PC 1** | **PC 2** |
| --- | --- | --- |
| %C | -0.214 | 0.601 |
| %N | -0.301 | 0.307 |
| C:N | 0.323 | 0.099 |
| Protein | 0.325 | -0.059 |
| WW | 0.300 | 0.313 |
| ORAC | 0.312 | -0.226 |
| TBARS | 0.324 | 0.070 |
| ORAC:TBARS | -0.324 | -0.078 |
| AChE | 0.217 | 0.595 |
| $\delta^{15}\text{N}$ | -0.324 | 0.078 |
| $\delta^{13}\text{C}$ | -0.323 | -0.103 |

The PC1 loadings also indicate higher values for both δ^15^N and δ^13^C; moreover, this increase was uncoupled from the oxidative status. The PC2 explaining the rest of the variance was mostly (>0.5) associated with positive loadings of %C and AChE. The projection of %C and AChE on the biplot indicated elevated %C in PH treatment and higher AChE activity in controls (Table 2).

### Mortality and exposure concentrations

Amphipod mortality was high in all treatments (51-77 %; Table S1, Supporting Information). It decreased with increasing propranolol concentration (GLM: p < 0.03) with significantly lower values in the PH treatment compared to the control amphipods (Tukey test: p < 0.02). The propranolol concentrations in the system were below the quantification limit in the control treatment, whereas the levels in the water were close to the nominal concentrations for both the PL and PH treatment: 108 ± 5.8 μg L^-1^ and 1058 ± 37 μg L^-1^, respectively (mean ± SE). Moreover, the concentrations measured in amphipods were twice as high in the PH treatment compared to PL, 6.3 and 3.2 μg g WW^-1^, respectively (Table S1).

### Hypothesis testing

#### H1

As hypothesized, the nitrogen content was significantly positively affected by propranolol concentration, with %N varying from 8.2% in PL and control to 8.6% in the PH treatment (Table 3; Figure 3A).

**Figure 2.**
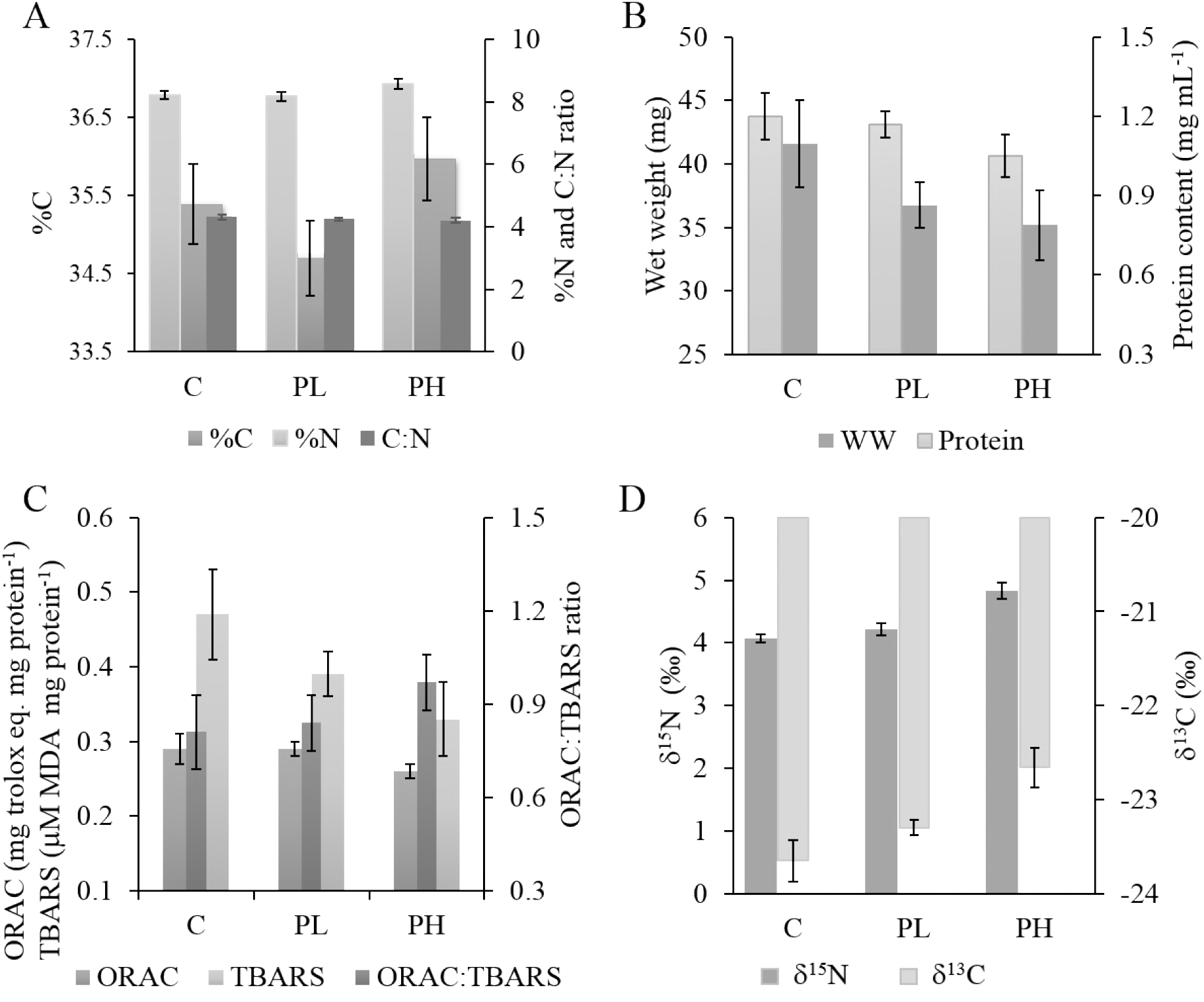
The elemental composition (%C and %N) and the C:N ratio in amphipods (A), body size and protein content (B), biomarkers of oxidative stress (antioxidant capacity assayed as ORAC; lipid peroxidation assayed as TBARS; balance between antioxidative and pro-oxidative activities assayed as the ORAC:TBARS ratio) (C), and stable isotope composition (δ^15^N and δ^13^C) (D) in amphipods exposed to propranolol. Control (0 μg L^-1^ propranolol), PL (100 μg L^-1^ propranolol), PH (1000 μg L^-1^ propranolol). Data are shown as group means and error bars represent min and max values.

**Table 3.** Generalized linear and generalized least square models testing treatment effects on %N, WW, TBARS, ORAC, AChE, δ^15^N, δ^13^C. Propranolol, concentration of propranolol (μg L^-1^); %N, nitrogen content; %C, carbon content; WW, wet weight; TBARS, thiobarbituric acid reactive substances as proxy for reactive oxygen species; ORAC, oxygen radical absorbance capacity as proxy for antioxidative capacity; AChE, acetylcholinesterase activity. When used as response variables, the values for WW, AChE, TBARSp, δ^15^N and δ^13^C were Box-Cox transformed.

| Dependent variable | Explanatory variables | Estimate | SE | t | p value |
| --- | --- | --- | --- | --- | --- |
| %N | propranolol | 0.00005 | 0.00002 | 2.055 | **0.046** |
| Protein | propranolol | -0.00014 | 0.00009 | -1.516 | 0.137 |
|  | %N | 0.282 | 0.100 | 2.821 | **0.007** |
| | propranolol × %N | -0.0004 | 0.0002 | -2.551 | **0.015** |
| WW | protein | 0.023 | 0.003 | 6.814 | **0.000** |
| TBARS | propranolol | -0.0003 | 0.0001 | -2.155 | **0.037** |
| ORAC | WW | 3.061 | 0.711 | 4.307 | **0.000** |
| AChE | propranolol | -0.006 | 0.002 | -2.408 | **0.020** |
|  | ORAC:TBARS | 0.080 | 0.018 | 4.494 | **0.000** |
| $\delta^{15}\text{N}$ | propranolol | 0.0002 | 0.00003 | 5.127 | **0.000** |
|  | ORAC:TBARS | 0.087 | 0.039 | 2.203 | **0.033** |
| $\delta^{13}\text{C}$ | propranolol | 0.001 | 0.0002 | 4.171 | **0.000** |
|  | %C | -0.151 | 0.051 | -2.958 | **0.005** |

#### H2

The protein content variability was significantly positively related to %N, with a significant negative *propranolol* × *%N* interaction effect, indicating that at higher propranolol concentration less nitrogen was allocated to proteins (Table 3).

#### H3

The amphipods from the PH treatment had the lowest WW; they were by 4% and 17% smaller compared to the PL and control groups, respectively (Figure 3B). The protein content was the best positive predictor of WW (Table 3).

#### H4-6

As hypothesized, the TBARS levels were negatively related to the propranolol concentration, with the lowest values observed in the PH treatment where were lower by 17% and 35% compared to those measured in the PL and control amphipods, respectively (Figure 3C). Also, a significant inhibition of AChE activity was observed in propranolol-exposed animals. The ORAC values were significantly positively related to WW; moreover, a significant positive relationship between AChE and the ORAC:TBARS ratio was found (Table 3).

#### H7-8

The δ^15^N values were significantly positively related to propranolol concentration and the ORAC:TBARS ratio (Table 3). Thus, the δ^15^N values were elevated in the animals exposed to propranolol and with the prevalence of the antioxidants. In PH treatment, the δ^15^N values were higher by 0.7 ‰, with mean values of 4.8 ‰ compared to 4.1-4.2 ‰ in PL and control groups (Figure 3D). As hypothesised, the δ^13^C values were best explained by propranolol concentration and %C as significant positive and negative predictors, respectively. The amphipod δ^13^C values in the PH treatment (-22.7 ‰) were elevated by ~1 ‰, compared to the PL (-23.3 ‰) and control (-23.7 ‰) animals (Figure 3D).

## Discussion

In animals, the majority of drugs either (i) mimic or inhibit normal physiological/biochemical processes; (ii) inhibit pathological processes; or (iii) inhibit vital processes of endo-or ectoparasites/microorganisms. Our study aimed to evaluate whether exposure to propranolol that belongs to the first group would cause predictable effects in crustaceans with regard to their stable isotope ratios and oxidative status. Based on the known targets and MOAs of propranolol, we expected to observe a lower scope for growth, improved oxidative balance, and AChE inhibition. As isotopic fractionation is a function of growth and metabolism [5], we expected to observe higher δ-values in the exposed animals. Most of the hypothesized effects (Table 1; Figure 1) were indeed observed, albeit only because of the magnitude of responses in the highest propranolol concentration (1058 μg L^-1^). The two exposure concentrations resulted in internal propranolol concentrations (3-6 μg g^-1^) that were within the range of the therapeutic concentrations reported in rabbit (~5 μg g^-1^) and human (0.47-11.67 μg g^-1^) brain, respectively [43]; thus, the exposure levels were consistent with the therapeutic doses for this drug.

As hypothesized (H1), the nitrogen content was positively related to propranolol concentration and in agreement with the increased nitrogen retention observed in propranolol-exposed rats [21]. The positive effect of %N on the protein content (H2) was not particularly surprising as nitrogen constitute about 15-17% of cell protein [44,45]. However, the significant propranolol × %N interaction effect on protein content indicates that with increasing dose, less nitrogen was allocated to proteins. The decrease in protein-bound nitrogen (and a concomitant increase in non-protein nitrogen, presumably as ammonia) is in line with the elevated nitrogen excretion induced by propranolol in humans [46]. Also, a part of the non-protein nitrogen could have been associated with free amino acids or shorter peptides, indicating either an elevated breakdown or slow synthesis of proteins [20]. It is also possible, however, that in the slow-growing animals, the relative contribution of the exoskeleton to body mass was higher compared to that in the amphipods growing faster, which would result in the overall decrease of the protein-bound nitrogen in the former.

The consequence of the propranolol-induced inhibition of protein synthesis is growth inhibition that was both biologically (up to 17%) and statistically significant (Table 3). The reduced growth could be a consequence of a reduced food intake related to behavioral change and slow feeding in the propranolol treatments as described for rabbits [47] and bivalves [48]. In these studies, the reduced growth rate was associated with reduced food intake as a result of the propranolol exposure and not related to the β-adrenoreceptor-blocking activity of the drug. Moreover, the inhibited AChE activity observed in the propranolol-exposed amphipods (this study) and mussels [48,49] is also a manifestation of the behavioural response contributing to the possible decline in motility and food acquisition [50,51]. Interestingly, the observed growth inhibition was not translated into higher mortality in the exposed animals.

On the contrary, the mortality was significantly lower in the highest propranolol concentration compared to the controls, suggesting that changes in energy budget exerted by the exposure were promoting lower biomass accumulation yet increased survivorship. Such trade-offs between individual growth and mortality are well-known in ecology and occur both inter-and intra-specifically as a part of general adaptation and fitness optimization processes [52].

The oxidative balance assayed as lipid peroxidation (TBARS) and ORAC:TBARS ratio was predicted to improve in the propranolol exposure (H4; Figure 3C). Indeed, the levels of TBARS were ~17% and 35% lower in PH compared to PL and the control, respectively, which is in line with reports showing an ameliorating effect of propranolol on lipid peroxidation [22,23]. However, no direct effects of propranolol on the antioxidant levels and no relationship between ORAC and TBARS (H5) were found. The amphipod body size was the single best predictor for ORAC, with smaller amphipods having lower ORAC levels. The opposite, i.e., a decrease in antioxidant capacity with increasing body size/age, has been reported for animals across multiple ontogenetic stages and broad size span [53,54]. In our experiment, however, the amphipods were within a relatively narrow size range (1.1 to 1.5 cm) and, thus, the ontogenetic component was not likely to be detectable. Instead, the smaller body size in the propranolol treatments may reflect a decrease in food consumption and concomitant intake of dietary antioxidants resulting in lower ORAC levels. Finally, accelerated growth itself is a pro-oxidative process that drives elevated antioxidant production [55], which implies that slower-growing individuals would have lower ORAC values.

Both propranolol exposure and the oxidative balance were significant predictors of the AChE inhibition. However, contrary to Hypothesis 6, the relationship between AChE and ORAC:TBARS ratio was positive (Table 3). In humans, the positive effect of moderate oxidative stress on AChE activity has been linked to hydrogen peroxide-mediated inactivation of the enzyme [56], with concomitant positive feedback on AChE expression. Similar feedback between AChE activity and oxidative status has previously been suggested for the amphipod *Monoporeia affinis* exposed to contaminated sediments and hypoxia [57]. The latter study also reported a significant positive relationship between AChE and the RNA:DNA ratio, which is in line with the observed decrease in physical activity and feeding rate in *Gammarus* in concert with AChE inhibition [51] and the observed propranolol effects on growth and AChE (this study). Therefore, propranolol-induced inhibition of AChE may, at least in part, explain effects attributed to the antagonistic effects of propranolol on the serotonin receptor [58]. For instance, the reduced motility and feeding rate due to neurological impairments could reduce both antioxidant intake and oxidative damage, thus corroborating the observed ORAC and TBARS responses. Taken together, these findings indicate that propranolol-induced AChE inhibition may play an important role in both sub-organismal and organism-level effects of propranolol exposure in crustaceans.

As expected (H7-H8), we found significant positive effects of propranolol on both δ^15^N and δ^13^C values, whereas body size was not a significant predictor of the isotopic signatures. In crustaceans exposed to organochlorine and PCBs, the effects on δ^15^N and δ^13^C values were related to growth and metabolic rates [8,59]. Consequently, the metabolic detoxification costs were implicated in altering energy budget, increasing turnover, and isotope fractionation. However, the exposure to propranolol does not have to increase energy expenditure and, therefore, the mechanisms for the shifts in isotopic fractionation may differ from those acting solely via growth and metabolic turnover. For example, these pathways could involve alterations in cAMP signaling induced by the antagonistic effect of propranolol on the 5-HT1 receptor, previously observed in bivalves [28] or stem from the indirect effects of propranolol on the muscarinic acetylcholine receptors [60]. Changes in metabolic pathways can alter bulk δ^15^N [61] in response to alterations in ^15^N fractionation pattern of specific amino acids and their relative abundances [62]. The observed increase in δ^15^N is, therefore, in line with the significant changes in amino acid concentrations observed in *Gammarus* exposed to propranolol [63]. This correlative evidence suggests that greater isotope fractionation was related to changes in protein metabolism induced by propranolol.

Also, a shift towards antioxidative processes was associated with higher δ^15^N values as suggested by the positive effect of the ORAC:TBARS ratio on the δ^15^N values. This is in line with the hypothsis that isotope-reinforced biomolecules may better resist oxidative damage as was observed in yeasts [16]. In another study with yeasts, heavy hydrogen isotope has been found to confer resistance to oxidative phosphorylation, the major source of cellular ROS [64]. Hence, the ^15^N-enriched tissues would then emerge as a consequence of selective reactions between ROS and isotopically light biomolecules or as an adaptation to consistently elevated ROS production.

The relationships between isotopic composition and oxidative status are complex, because many dietary and non-dietary factors may affect both isotope markers and biomarkers of oxidative status. In field studies, effects of diet and physiological status on the isotopic signatures are particularly difficult to disentangle, because dietary sources with great differences in their isotopic signatures would mask the isotope effect on the oxidative status and vice versa. For example, a positive relationship between δ^15^N and oxidative damage in Gentoo penguins [10] was related to a combination of varying intake of antioxidants and foraging efforts associated with oxidative costs. Therefore, laboratory studies are particularly important for understanding the underlying causes of the observed isotope signatures in field studies. Since the amphipods in our study were all presented with the same food options and kept in the confined, controlled environment, no effects of variation in antioxidants for different food sources or differences in specific foraging costs as well as environmental factors may have confounded the isotopic signatures.

In summary, we have shown that the effects of the pharmaceutical propranolol on both stable isotope values and oxidative stress were indeed predictable in *Gammarus.* More studies are needed to investigate how MOA might be used to predict biomarker, nutrient allocation, and isotopic changes in non-target organisms possessing evolutionarily conserved targets for this drug, and, most probably, other pharmaceuticals with similar effect pathways. Moreover, a positive link between δ^15^N values and the oxidative balance was predicted and confirmed to exist in the exposed animals, which can be used for interpretation of biomarkers of oxidative stress and stable isotope data in ecotoxicological and ecological surveys. The fact that isotopic signal can be confounded by non-dietary parameters, including environmental stress, is also of high relevance for food web reconstructions based on stable isotope signatures, particularly, in environments chronically exposed to environmental contaminants, including pharmaceuticals.

## Supporting information

SI includes information on the observed mortality and quantifications of propranolol in amphipod and water samples (Table S1), statistical evaluation of the aquarium effects on the measured endpoints (Table S2), a summary of the measured endpoints (Table S3) and a diagram illustrating effects induced by propranolol (Figure S1).

## Acknowledgments

This study was supported by the Isotope ecology network in the Baltic Sea region (Swedish Institute, Sweden), State Key Laboratory of Pollution Control and Resource Reuse Foundation (Tongji University) (No. PCRRIC16003), and by the Swedish Research Council (project number 639-2013-6913).

## References

1. McCue MD. Tracking the Oxidative and Nonoxidative Fates of Isotopically Labeled Nutrients in Animals. BioScience. 2011;61: 217–230. doi:10.1525/bio.2011.61.3.7

2. Reitsema LJ. Beyond diet reconstruction: Stable isotope applications to human physiology, health, and nutrition: Stable Isotopes and Physiology. American Journal of Human Biology. 2013;25: 445–456. doi:10.1002/ajhb.22398

3. Carravieri A, Bustamante P, Churlaud C, Fromant A, Cherel Y. Moulting patterns drive within-individual variations of stable isotopes and mercury in seabird body feathers: implications for monitoring of the marine environment. Mar Biol. 2014;161: 963–968. doi:10.1007/s00227- 014-2394-x

4. Martinez del Rio C, Wolf BO. Mass-balance models for animal isotopic ecology. In: Starck MJ, Wang T, editors. Physiological and Ecological Adaptations to Feeding in Vertebrates. Enfield, NH: Science Publishers; 2005. pp. 141–174.

5. Gorokhova E. Individual growth as a non-dietary determinant of the isotopic niche metrics. Kurle C, editor. Methods in Ecology and Evolution. 2017; doi:10.1111/2041-210X.12887

6. Robbins CT, Felicetti LA, Sponheimer M. The effect of dietary protein quality on nitrogen isotope discrimination in mammals and birds. Oecologia. 2005;144: 534–540. doi:10.1007/s00442-005-0021-8

7. Power M, Guiguer KRRA, Barton DR. Effects of temperature on isotopic enrichment in Daphnia magna: implications for aquatic food-web studies. Rapid Commun Mass Spectrom. 2003;17: 1619–1625. doi:10.1002/rcm.1094

8. Ek C, Karlson AML, Hansson S, Garbaras A, Gorokhova E. Stable Isotope Composition in Daphnia Is Modulated by Growth, Temperature, and Toxic Exposure: Implications for Trophic Magnification Factor Assessment. Environmental Science & Technology. 2015;49: 6934–6942. doi:10.1021/acs.est.5b00270

9. Shaw-Allen PL, Romanek CS, Bryan AL, Brant H, Jagoe CH. Shifts in relative tissue delta15N values in snowy egret nestlings with dietary mercury exposure: a marker for increased protein degradation. Environ Sci Technol. 2005;39: 4226–4233.

10. Beaulieu M, González-Acuña D, Thierry A-M, Polito MJ. Relationships between isotopic values and oxidative status: insights from populations of gentoo penguins. Oecologia. 2015;177: 1211–1220. doi:10.1007/s00442-015-3267-9

11. Hoefs J, editor. Isotope Fractionation Processes of Selected Elements. Stable Isotope Geochemistry. Berlin, Heidelberg: Springer Berlin Heidelberg; 2009. pp. 35–92. doi:10.1007/978-3-540-70708-0_2

12. Livingstone DR. Contaminant-stimulated reactive oxygen species production and oxidative damage in aquatic organisms. Mar Pollut Bull. 2001;42: 656–666.

13. Furuhagen S, Liewenborg B, Breitholtz M, Gorokhova E. Feeding Activity and Xenobiotics Modulate Oxidative Status in Daphnia magna: Implications for Ecotoxicological Testing. Environ Sci Technol. 2014;48: 12886–12892. doi:10.1021/es5044722

14. Alonso-Alvarez C, Bertrand S, Devevey G, Prost J, Faivre B, Sorci G. Increased susceptibility to oxidative stress as a proximate cost of reproduction. Ecology Letters. 2004;7: 363–368. doi:10.1111/j.1461-0248.2004.00594.x

15. Shchepinov MS. Reactive oxygen species, isotope effect, essential nutrients, and enhanced longevity. Rejuvenation Res. 2007; 10: 47–59. doi:10.1089/rej.2006.0506

16. Hill S, Hirano K, Shmanai VV, Marbois BN, Vidovic D, Bekish AV, et al. Isotope-reinforced polyunsaturated fatty acids protect yeast cells from oxidative stress. Free Radic Biol Med. 2011;50: 130–138. doi:10.1016/j.freeradbiomed.2010.10.690

17. Banas D, Vollaire Y, Danger M, Thomas M, Oliveira-Ribeiro CA, Roche H, et al. Can we use stable isotopes for ecotoxicological studies? Effect of DDT on isotopic fractionation in Perca fluviatilis. Chemosphere. 2009;76: 734–739. doi:10.1016/j.chemosphere.2009.05.033

18. Ek C, Holmstrand H, Mustajärvi L, Garbaras A, Bariseviciute R, Sapolaite J, et al. Using compound-specific and bulk stable isotope analysis for trophic positioning of bivalves in contaminated Baltic Sea sediments. Environ Sci Technol. 2018;52: 4861–4868. doi:10.1021/acs.est.7b05782

19. Gunnarsson L, Jauhiainen A, Kristiansson E, Nerman O, Larsson DGJ. Evolutionary Conservation of Human Drug Targets in Organisms used for Environmental Risk Assessments. Environ Sci Technol. 2008;42: 5807–5813. doi:10.1021/es8005173

20. Kwiatkowska-Patzer B, Zalewska T. Effect of propranolol upon protein and proteolytic synthesis activity in hypertrophic myocardium. Basic Res Cardiol. 1988;83: 43–47.

21. Dickerson RN, Fried RC, Bailey PM, Stein TP, Mullen JL, Buzby GP. Effect of propranolol on nitrogen and energy metabolism in sepsis. J Surg Res. 1990;48: 38–41.

22. Kramer JH, Murthi SB, Wise RM, Mak I-T, Weglicki WB. Antioxidant and lysosomotropic properties of acute D-propranolol underlies its cardioprotection of postischemic hearts from moderate iron-overloaded rats. Exp Biol Med (Maywood). 2006;231: 473–484.

23. Kramer JH, Spurney CF, Iantorno M, Tziros C, Chmielinska JJ, Mak IT, et al. d-Propranolol protects against oxidative stress and progressive cardiac dysfunction in iron overloaded rats. Can J Physiol Pharmacol. 2012;90: 1257–1268. doi:10.1139/y2012-091

24. Wilkens JL. The control of cardiac rhythmicity and of blood distribution in crustaceans. Comparative Biochemistry and Physiology Part A: Molecular & Integrative Physiology. 1999;124: 531–538. doi:10.1016/S1095-6433(99)00146-4

25. Alexander BS, Wood MD. Stereoselective blockade of central [3H]5-hydroxytryptamine binding to multiple sites (5-HT1A, 5-HT1B and 5-HT1C) by mianserin and propranolol. J Pharm Pharmacol. 1987;39: 664–666.

26. Tierney AJ. Structure and function of invertebrate 5-HT receptors: a review. Comp Biochem Physiol, Part A Mol Integr Physiol. 2001; 128: 791–804.

27. Covi JA, Chang ES, Mykles DL. Conserved role of cyclic nucleotides in the regulation of ecdysteroidogenesis by the crustacean molting gland. Comp Biochem Physiol, Part A Mol Integr Physiol. 2009;152: 470–477. doi:10.1016/j.cbpa.2008.12.005

28. Franzellitti S, Buratti S, Valbonesi P, Capuzzo A, Fabbri E. The β-blocker propranolol affects cAMP-dependent signaling and induces the stress response in Mediterranean mussels, Mytilus galloprovincialis. Aquat Toxicol. 2011;101: 299–308. doi:10.1016/j.aquatox.2010.11.001

29. Alkondon M, Ray A, Sen P. Tissue cholinesterase inhibition by propranolol and related drugs. J Pharm Pharmacol. 1986;38: 848–850.

30. Dzialowski EM, Turner PK, Brooks BW. Physiological and reproductive effects of beta adrenergic receptor antagonists in Daphnia magna. Arch Environ Contam Toxicol. 2006;50: 503–510. doi:10.1007/s00244-005-0121-9

31. Wiklund A-KE, Oskarsson H, Thorsén G, Kumblad L. Behavioural and physiological responses to -pharmaceutical exposure in macroalgae and -grazers from a Baltic Sea littoral community. Aquatic Biology. 2011;14: 29–39. doi:10.3354/ab00380

32. Ternes TA. Occurrence of drugs in German sewage treatment plants and rivers. Water Research. 1998;32: 3245–3260. doi:10.1016/S0043-1354(98)00099-2

33. Huggett DB, Khan IA, Foran CM, Schlenk D. Determination of beta-adrenergic receptor blocking pharmaceuticals in united states wastewater effluent. Environmental Pollution. 2003;121: 199–205. doi:10.1016/S0269-7491(02)00226-9

34. Oskarsson H, Wiklund A-KE, Thorsén G, Danielsson G, Kumblad L. Community Interactions Modify the Effects of Pharmaceutical Exposure: A Microcosm Study on Responses to Propranolol in Baltic Sea Coastal Organisms. PLOS ONE. 2014;9: e93774. doi:10.1371/journal.pone.0093774

35. Ou B, Hampsch-Woodill M, Prior RL. Development and Validation of an Improved Oxygen Radical Absorbance Capacity Assay Using Fluorescein as the Fluorescent Probe. J Agric Food Chem. 2001;49: 4619–4626. doi:10.1021/jf010586o

36. Rikans LE, Hornbrook KR. Lipid peroxidation, antioxidant protection and aging. Biochimica et Biophysica Acta (BBA) - Molecular Basis of Disease. 1997;1362: 116–127. doi:10.1016/S0925- 4439(97)00067-7

37. Vehmaa A, Hogfors H, Gorokhova E, Brutemark A, Holmborn T, Engström-Öst J. Projected marine climate change: effects on copepod oxidative status and reproduction. Ecol Evol. 2013;3: 4548–4557. doi:10.1002/ece3.839

38. Rickwood CJ, Galloway TS. Acetylcholinesterase inhibition as a biomarker of adverse effect. Aquatic Toxicology. 2004;67: 45–56. doi:10.1016/j.aquatox.2003.11.004

39. Smith PK, Krohn RI, Hermanson GT, Mallia AK, Gartner FH, Provenzano MD, et al. Measurement of protein using bicinchoninic acid. Anal Biochem. 1985;150: 76–85.

40. Hodges DM, DeLong JM, Forney CF, Prange RK. Improving the thiobarbituric acid-reactive- substances assay for estimating lipid peroxidation in plant tissues containing anthocyanin and other interfering compounds. Planta. 1999;207: 604–611. doi:10.1007/s004250050524

41. Ellman G, Courtney K, Andres V, Featherstone R. A new and rapid colorimetric determination of acetylcholinesterase activity. Biochemistry and Pharmacology. 1961;7: 88–95.

42. Hammer Ø, Harper DA, Ryan PD. PAST: Paleontological Statistics Software Package for Education and Data Analysis. Palaeontologia Electronica. 2001;4: 1–9.

43. Myers MG, Lewis PJ, Reid JL, Dollery CT. Brain concentration of propranolol in relation to hypotensive effect in the rabbit with observations on brain propranolol levels in man. J Pharmacol Exp Ther. 1975;192: 327–335.

44. Winberg GG. Methods for the estimation of production of aquatic animals. Academic 882 Press Inc, London, United Kingdom; 1971.

45. Elser JJ, Sterner RW, Gorokhova E, and Fagan WF, Markow TA, Cotner JB, et al. Biological stoichiometry from genes to ecosystems. Ecology Letters. 2000;3: 540–550.

46. Acheson KJ, Ravussin E, Schoeller DA, Christin L, Bourquin L, Baertschi P, et al. Two-week stimulation or blockade of the sympathetic nervous system in man: influence on body weight, body composition, and twenty four-hour energy expenditure. Metab Clin Exp. 1988;37: 91–98.

47. Evemy KL, Cummins P, Littler WA. The Effect of Prolonged Administration of Propranolol and Timolol on the Growth and the Heart of Growing Rabbits. Clinical Science. 1981;60: 33–40. doi:10.1042/cs0600033

48. Solé M, Shaw JP, Frickers PE, Readman JW, Hutchinson TH. Effects on feeding rate and biomarker responses of marine mussels experimentally exposed to propranolol and acetaminophen. Analytical and Bioanalytical Chemistry. 2010;396: 649–656. doi:10.1007/s00216-009-3182-1

49. Singh AK, Spassova D. Effects of hexamethonium, phenothiazines, propranolol and ephedrine on acetylcholinesterase carbamylation by physostigmine, aldicarb and carbaryl: interaction between the active site and the functionally distinct peripheral sites in acetylcholinesterase. Comp Biochem Physiol C, Pharmacol Toxicol Endocrinol. 1998;119: 97–105.

50. Pavlov DD, Chuiko GM, Gerassimov YV, Tonkopiy VD. Feeding behavior and brain acetylcholinesterase activity in bream (Abramis brama L.) as affected by DDVP, an organophosphorus insecticide. Comp Biochem Physiol C, Comp Pharmacol Toxicol. 1992;103: 563–568.

51. Xuereb B, Lefèvre E, Garric J, Geffard O. Acetylcholinesterase activity in Gammarus fossarum (Crustacea Amphipoda): linking AChE inhibition and behavioural alteration. Aquat Toxicol. 2009;94: 114–122. doi:10.1016/j.aquatox.2009.06.010

52. Willmer P, Stone G, Johnston IA. Environmental physiology of animals. 2nd ed. Malden, Mass: Blackwell Pub; 2005.

53. Correia AD, Costa MH, Luis OJ, Livingstone DR. Age-related changes in antioxidant enzyme activities, fatty acid composition and lipid peroxidation in whole body Gammarus locusta (Crustacea: Amphipoda). Journal of Experimental Marine Biology and Ecology. 2003;289: 83–101. doi:10.1016/S0022-0981(03)00040-6

54. Barata C, Navarro JC, Varo I, Riva MC, Arun S, Porte C. Changes in antioxidant enzyme activities, fatty acid composition and lipid peroxidation in Daphnia magna during the aging process. Comp Biochem Physiol B, Biochem Mol Biol. 2005;140: 81–90. doi:10.1016/j.cbpc.2004.09.025

55. Speakman JR, Mitchell SE. Caloric restriction. Molecular Aspects of Medicine. 2011;32: 159–221. doi:10.1016/j.mam.2011.07.001

56. Schallreuter KU, Elwary SMA, Gibbons NCJ, Rokos H, Wood JM. Activation/deactivation of acetylcholinesterase by H2O2: more evidence for oxidative stress in vitiligo. Biochem Biophys Res Commun. 2004;315: 502–508. doi:10.1016/j.bbrc.2004.01.082

57. Gorokhova E, Löf M, Reutgard M, Lindström M, Sundelin B. Exposure to contaminants exacerbates oxidative stress in amphipod Monoporeia affinis subjected to fluctuating hypoxia. Aquatic Toxicology. 2013;127: 46–53. doi:10.1016/j.aquatox.2012.01.022

58. Huggett DB, Brooks BW, Peterson B, Foran CM, Schlenk D. Toxicity of select beta adrenergic receptor-blocking pharmaceuticals (B-blockers) on aquatic organisms. Arch Environ Contam Toxicol. 2002;43: 229–235. doi:10.1007/s00244-002-1182-7

59. Ek C, Gerdes Z, Garbaras A, Adolfsson-Erici M, Gorokhova E. Growth Retardation and Altered Isotope Composition As Delayed Effects of PCB Exposure in Daphnia magna. Environmental Science & Technology. 2016;50: 8296–8304. doi:10.1021/acs.est.6b01731

60. Hannan F, Hall LM. Muscarinic acetylcholine receptors in invertebrates: Comparisons with homologous receptors from vertebrates. In: Pichon Y, editor. Comparative Molecular Neurobiology. Basel: Birkhäuser Basel; 1993. pp. 98–145. doi:10.1007/978-3-0348-7265-2_6

61. Poupin N, Mariotti F, Huneau J-F, Hermier D, Fouillet H. Natural Isotopic Signatures of Variations in Body Nitrogen Fluxes: A Compartmental Model Analysis. PLOS Computational Biology. 2014;10: e1003865. doi:10.1371/journal.pcbi.1003865

62. Schmidt K, McClelland JW, Mente E, Montoya JP, Atkinson A, Voss M. Trophic-level interpretation based on 515N values: implications of tissue-specific fractionation and amino acid composition. Marine Ecology Progress Series. 2004;266: 43–58. doi:10.3354/meps266043

63. Gómez-Canela C, Miller TH, Bury NR, Tauler R, Barron LP. Targeted metabolomics of Gammarus pulex following controlled exposures to selected pharmaceuticals in water. Science of The Total Environment. 2016;562: 777–788. doi:10.1016/j.scitotenv.2016.03.181

64. Li X, Snyder MP. Can heavy isotopes increase lifespan? Studies of relative abundance in various organisms reveal chemical perspectives on aging. BioEssays. 2016;38: 1093–1101. doi:10.1002/bies.201600040

